# LED light gradient as a screening tool for light quality responses in model plant species

**DOI:** 10.1101/2020.10.08.320002

**Authors:** P. Lejeune, A. Fratamico, F. Bouché, S. Huerga Fernández, P. Tocquin, C. Périlleux

## Abstract

Current developments in light-emitting diodes (LEDs) technologies have opened new perspectives for sustainable and highly efficient indoor cultivation. The introduction of LEDs not only allows a reduction in the production costs on a quantitative level, it also offers opportunities to manipulate and optimise qualitative traits. Indeed, while plants respond strongest to red and blue lights for photosynthesis, the whole light spectrum has an effect on plant shape, development, and chemical composition. In order to evaluate LEDs as an alternative to traditional lighting sources, the species-specific plant responses to distinct wavelengths need to be evaluated under controlled conditions. Here, we tested the possibility to use light composition gradients in combination with semi-automated phenotyping to rapidly explore the phenotypic responses of different species to variations in the light spectrum provided by LED sources. Plants of seven different species (*Arabidopsis thaliana, Ocimum basilicum, Solanum lycopersicum, Brachypodium distachyon, Oryza sativa, Euphorbia peplus, Setaria viridis*) were grown under standard white fluorescent light for 30 days, then transferred to a Red:Blue gradient for another 30 days and finally returned to white light. In all species, differences in terms of dimension, shape, and color were rapidly observed across the gradient and the overall response was widely species-dependent. The experiment yielded large amounts of imaging-based phenotypic data and we suggest simple data analysis methods to aggregate the results and facilitate comparisons between species. Similar experimental setups will help achieve rapid environmental optimization, screen new crop species and genotypes, or develop new gene discovery strategies.

## 1. Introduction

Plants are sessile organisms that must rely on environmental cues to adapt their physiology and morphology to prevailing and changing conditions. Among those environmental cues, light is one of the most useful signals for plants. Not only does it fuel growth through photosynthesis, but it also brings information about the time of the day, the season, the surrounding environment, or the atmospheric conditions (1–4).

Light is perceived by photosynthetic pigments and by dedicated chromoproteins, called photoreceptors. In *Arabidopsis thaliana*, each of the five known photoreceptor families is sensitive to a specific region of the light spectrum, ranging from UV-B to near infrared (5). Through this complex sensing machinery, light quality controls multiple plant developmental processes, such as germination, growth under competing canopies, root development, and flowering (6–8). Photoreceptors are integrative triggers ensuring a fine-tuned response to the whole light spectrum (9, 10), while also interacting with hormonal pathways to coordinate plant growth and development (11, 12). Moreover, there is an interplay between the signaling function of light, which is efficient even at very low irradiances, and its energetic function in photosynthesis, since some of the responses triggered by photoreceptors have a direct impact on photosynthesis efficiency (leaf inclination, leaf flattening, chloroplast movement), carbon metabolism, biomass production, and stress responses (13–17).

In semi- or fully-controlled production environments, such as greenhouses or indoor farms, light is a limiting factor for crop and fruit yields. The use of supplemental artificial lighting is thus necessary in northern regions, especially in winter when shorter photoperiods and lower light intensities severely impact productivity (18). Moreover, with the continuous growth of the world population, artificial lighting is increasingly needed to support the growing demand for local food production in the emerging indoor urban farming infrastructures (19). During the last two decades, the improvement in the efficiency of light-emitting diodes (LEDs) has been the main driver in the development of these plant factories (20, 21). Given the high rate at which their luminous efficiency increases and their cost decreases, LEDs should soon outperform all other technologies for providing supplemental lighting in greenhouses (18, 22).

LEDs were invented in the 1960s and the range of available wavelengths has grown steadily since. The red and blue LEDs were the first whose efficiencies were sufficient for horticultural applications, and the fact that these wavelengths are the most efficient for photosynthesis obviously facilitated their adoption (23, 24). It was often shown that photosynthesis and growth benefit from a high Red:Blue ratio (17, 25–27), as expected from their respective quantum yield (28). However, thanks to the increase in available LED wavelengths, further studies revealed very complex responses to variations in the light spectrum. For instance, green and far-red wavelengths, which were initially neglected because of their low contribution to the action spectrum of photosynthesis, were shown to have a stimulating effect on photosynthesis in some conditions and could thus be useful to fine-tune crop and fruit productions (29–31). Moreover, because LED-based lightings enable the creation of “light recipes” by mixing and modulating an increasing number of available wavelengths, the trend is now to develop smart lighting applications (32). The goals are not only to fine-tune photosynthesis, growth and yield more efficiently, but also to improve the quality of crops by manipulating their secondary metabolism (33–36).

Given the extremely complex and species-dependent nature of light responses, comparing discrete experimental conditions would restrict the exploratory field and limit the significance of the results. Here, we screened the phenotypic responses of a panel of species to a Red:Blue gradient in order to maximize our understanding of the effect of varying ratios of these wavelengths across flowering plants. We chose to characterize seven model plants, based on their scientific and economical importance as well as their botanical diversity. We selected four dicot species: *Arabidopsis thaliana* (Brassicaceae), an obvious choice due to its popularity in academic research and the wealth of genomic and phenomic knowledge, *Solanum lycopersicum* (Solanaceae) and *Ocimum basilicum* (Lamiaceae), two interesting models for horticultural applications, as well as *Euphorbia peplus* (Euphorbiaceae), a wild species studied for its medicinal properties. We also selected three monocot model species (Poaceae): one tropical crop, *Oryza sativa*, one temperate species, *Brachypodium distachyon*, and finally one C4 wild species that is increasingly used in fundamental research, *Setaria viridis*.

Efficient phenotyping is another bottleneck for implementing LEDs into crop management and breeding applications. Over the last two decades, numerous publications have described novel phenotyping approaches to suit ever increasing fields of application. Technologies were developed to adjust the level of desired throughput, diversity, and scale of measured traits (i.e. cell, organ, plant, and canopy levels), to adapt to the plant growth facilities (e.g. field, greenhouse, indoor cabinets), and to serve various experimental aims (e.g. genomics, breeding, precision agriculture, screening of chemicals or bioactive compounds). These aspects have been extensively discussed in recent reviews (37–39). Imaging-based systems, thanks to their non-invasiveness and amenability to automation, have been increasingly used to measure plant traits since the late 1990s (40, 41), enabling the rapid collection of phenotypic data from larger populations of plants and at lower cost compared to manual approaches. Numerous variations of digital imaging setups have been developed with success to tackle a variety of applications and scientific questions (42). The implementation of a phenotyping pipeline implies numerous and inevitable compromises between the scope, the desired quality, the timelines, and the available budget. Commercial ready-to-use solutions are available for high-throughput, high-resolution, highly automated imaging platforms but they are still expensive due to the niche market and the high degree of customization. However, it is possible to construct simple low-cost imaging stations with sufficient image quality and speed, using off-the-shelf electro-mechanics, cameras, software, and open-source analysis tools (43–45). Here, we assembled an in-house, simple, and cost-efficient RGB imaging setup in order to capture basic but precise and reproducible biometrics (e.g. plant dimensions, shape factors, color indices) that enabled us to quantify the effects of LED lighting on the phenotype of selected plant species.

## 2. Materials and Methods

### 2.1. Experimental and Technical Design

Figure 1 summarizes the experimental workflow. Seedlings were first grown for 30 days in small Jiffypots^®^ under “normal” white light before being transplanted in standard 12-cm pots and transferred under a Red:Blue gradient. The purpose of starting the cultivation under white light was to avoid mixing the effects of light quality on germination and seedling establishment with its effects on later growth. After 30 days under the gradient, plants were re-transferred to white light. For each species, a group of plants was kept continuously under white light as a control. The light spectrum was recorded for each individual plant under the gradient conditions and white light. The Red:Blue ratio (PFD_Red_(600-700nm) over PFD_Blue_(400-500nm)) was calculated for individual plants based on the spectral light measurements performed at each plant position at the beginning of the gradient treatment. Plants were imaged every 3-4 days during the gradient treatment and after return to white light. Three types of phenotypic measurements were derived from the images: dimensions, shape factors, and color indices. These measurements were used to estimate the variation of plant size, morphology, and pigmentation along the Red:Blue gradient and across time. The leaf chlorophyll content was measured at the end of the gradient treatment. Data processing and analysis followed as described in section 2.6.

**Figure 1.**
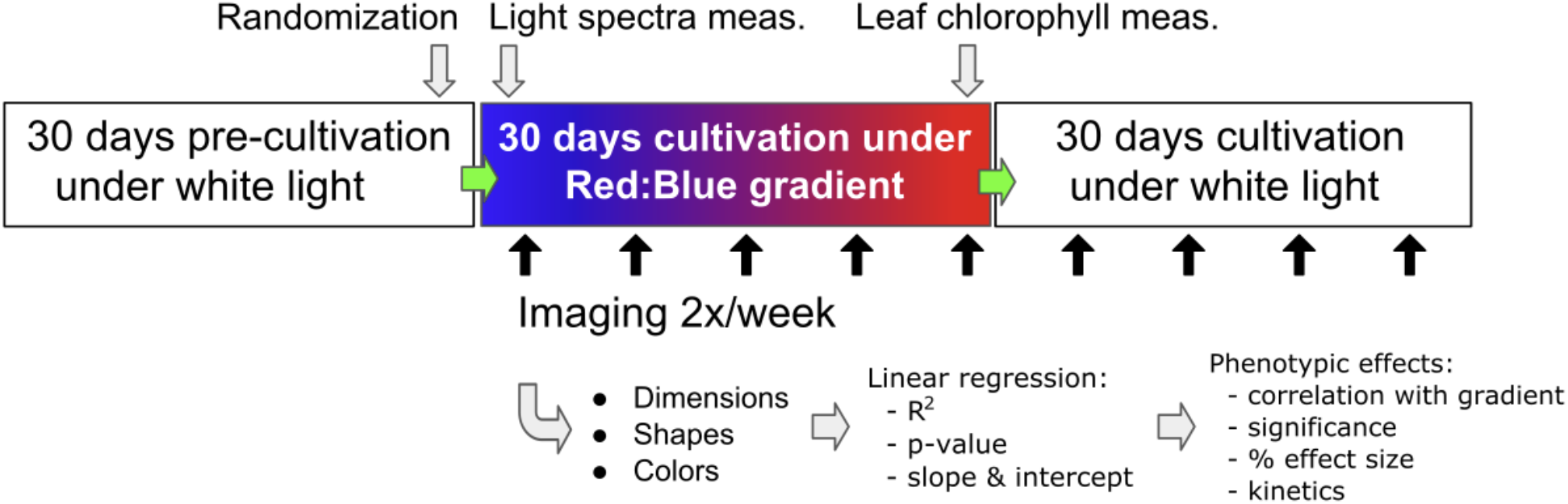
Experimental workflow.

### 2.2. Plant materials

*Arabidopsis thaliana* Col-0 seeds were obtained from a public seedbank (NASC, Nottingham, UK) and *Brachypodium distachyon* Bd21-3 seeds from Prof. R. Amasino (University of Wisconsin, USA). Seeds of *Euphorbia peplus* were obtained from fairdinkumseeds.com (Queensland, Australia). Seeds of *Setaria viridis* A10.1 were obtained from the USDA Iowa State University Agricultural Research Service (Ames, IO, USA). Seeds of *Solanum lycopersicum* cv. Ailsa Craig were obtained from TGRC (Davis, CA, USA). *Ocimum basilicum* cv. Genovese seeds were obtained from Le Jardin de Bellecourt (Bellecourt, Belgium). Seeds of *Oryza sativa* cv. Nipponbare were obtained from IRRI (Los Baños, Laguna, The Philippines).

### 2.3. Growth conditions

#### Germination

Seeds were sown in 4.5 cm fiber pots (Jiffypots^®^, Jiffy, Zwijndrecht, The Netherlands) filled with a 4:1 (vol:vol) mix of leaf mould and baked clay granules. The fiber pots were placed on 120 x 18 x 14 cm cultivation gutters (Goponic, Nouméa, France) and irrigated by capillarity through a wet cultivation felt mat (Feutriplanta^®^, Jardirama, Warsage, Belgium). The felt mat was kept continuously moist with felt wicks dipping in the water through holes (one every 10 cm) in the decks of the gutters (Figure 2). This capillarity system provides “on-demand” irrigation and avoids water excess or substrate compaction problems. The gutters were placed for 30 days in a Conviron PGV36 growth room (Conviron, Winnipeg, Canada) at 21°C day/night, 70% relative humidity, 12-h photoperiod, at an irradiance of ± 130-150 μE.m^−2^.s^−1^ provided by Sylvania Luxline Plus T5 FHO 54W tubes (Osram-Sylvania, Wilmington, MA, USA) delivering 4000K white light. Depending on the species, germination started between 1 and 2 weeks after sowing.

**Figure 2.**
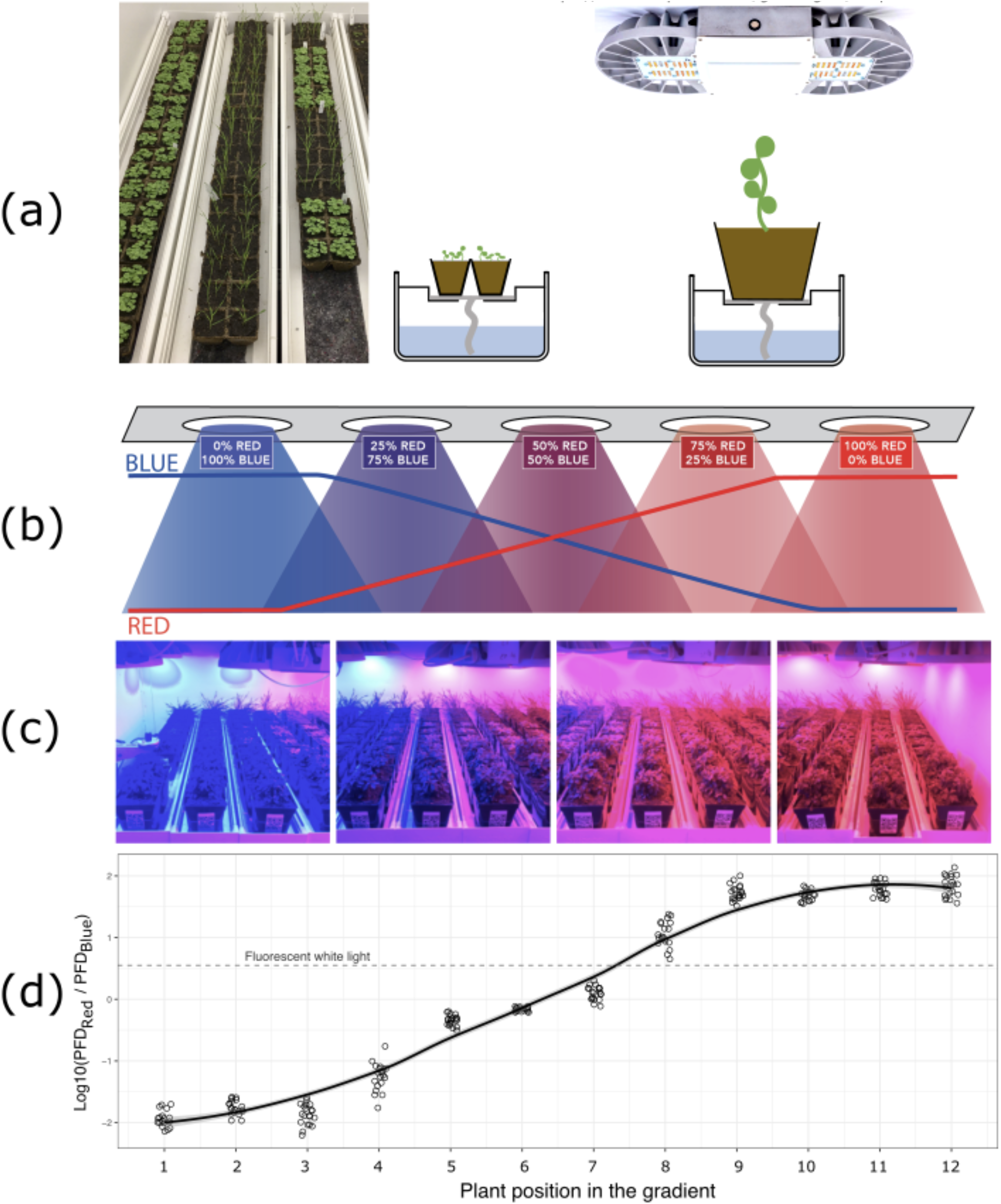
Experimental cultivation setup under the Red:Blue light gradient. (a) Left: 30-day-old plantlets growing in 4.5 cm fiber pots (Jiffypots) inside cultivation gutters lined with felt mats and wicks absorbing water. Right: cultivation system after transplantation in 12-cm pots under the Lumiatec LED luminaries. (b) Red:Blue gradient setup. Arrangement and setting of the 5 clusters of 3 LED luminaries in the phytotronic cabinet. (c) View of the phytotronic cabinet during the experiment. (d) Red:Blue ratio measured at each plant position. PFD = Photon Flux Density. Picture of the luminary in (a) is a courtesy of Araponics (Liège, Belium, https://www.araponics.com/grow-lights/63-phs16.html).

#### Plant Growth

After the initial 30 days under white light, Jiffypots^®^ with weak or abnormal plantlets were discarded and the others were transplanted into 12-cm square plastic cultivation pots filled with 1.5 L of leaf mould and baked clay (4:1) mixed with 6 gr.L^−1^ of slow release fertilizer (Osmocote Exact Standard 5-6 M, ICL Specialty Fertilizers). The pots were fitted at the bottom with a 2 x 10 cm felt wick and randomly placed on the deck of the cultivation gutters described above. The gutters were placed in Conviron PGV36 growth rooms under the same conditions than during germination, except for the lighting which was provided either by white fluorescent tubes (same type as above) or by adjustable 16 channels LED luminaries (described below). Each room had a 1.3 x 2.4 m (3 m^2^) cultivation area, allowing 12 gutters of 10 pots. The placement of the plants was organized in rows and columns so that each pot could be registered by Room:Row:Column coordinates and labelled with a unique QR-code. We used three contiguous rows per species, except for *Arabidopsis* that had four rows. A randomization step was performed for each species within the block of three rows to avoid any bias while placing the transplanted pots in the rooms. Right after transplantation, we only kept one plant per pot, except for *E. peplus* (6 plants/pot) and *O. basilicum* (up to 9 plants/pot) to account for their usual mode of cultivation in bushes.

After transplantation (day 30), the plants were taken out of the growth rooms twice a week for imaging and placed back at the same location. On day 60, all plants were transferred to white light conditions, grown and imaged for at least two more weeks. Beyond that point, plants of a given species were discarded if more than 50% were showing signs of flowering. After 30 days under white light, the experiment was stopped.

### 2.4 Spectrally adjustable LED lightings

Three phytotronic cabinets were equipped with 15 Lumiatec PHS:: 16 (300W) luminaries (GDTech, Alleur, Belgium) each. These luminaries are controllable over 16 independent channels (2x blue 455 nm, 6x white 4000K, 1x green 520 nm, 1x yellow 593 nm, 2x red 635 nm, 2x hi-red 660 nm, 1x far-red 730 nm, and 1x UV 280 nm) of 6 LEDs each. The 15 luminaries were regularly distributed as a 5 x 3 pattern at a distance of 45 cm between each other in order to guarantee optimal spectral homogeneity in the 3 m^2^ culture area (Figure 2). The luminaries were controlled per clusters of 3 using the Lumiatec control interface and the Blue and Red channels were adjusted as shown in Figure 2b in order to create a gradient of Red:Blue ratio (Figure 2d). The light spectrum and intensity across the growth chambers were monitored using a HiPoint HR-550 spectrophotometer (TAIWAN HIPOINT CORP., Kaohsiung, Taiwan).

### 2.5 Phenotyping platform, imaging process, and data analysis

Phenotypic data were collected twice a week using an in-house imaging cabinet (Figure 3). The setup was built with aluminum profiles supporting white PVC walls. The cabinet was illuminated by 25 x 25 cm white light LED panels (Araponics, Liège, Belgium). Lighting was optimised for taking pictures with a diffusive back-lit white background for side-view images and a black cloth background for top-view images. A step-motor platform was used to rotate the plants while two CMOS RGB 12 Mpx industrial cameras (Dalsa Genie-nano 4040, Dalsa, Waterloo, Canada) acquired plant images and one color HD webcam (Logitech, Lausanne, Switzerland) read QR-coded labels on the pots. The Genie-Nano cameras were fitted with high resolution 25-mm focal length Tamron M111FM25 lenses, which allowed to image plants up to 150-cm high and 100-cm wide with an estimated smallest detail size of +/− 0.5 mm at a working distance of 200 cm, based on sensor dimensions (14.2 x 10.4 mm, 4112 x 3008 pixels) and lens optical resolution (3.1 μm “pixel pitch”). Diaphragm closure of the lenses was set to F8.0, exposure time to 0.2 msec and gain to 6. A blueprint of the imaging setup is provided as supplementary material (Figure S1).

**Figure 3.**
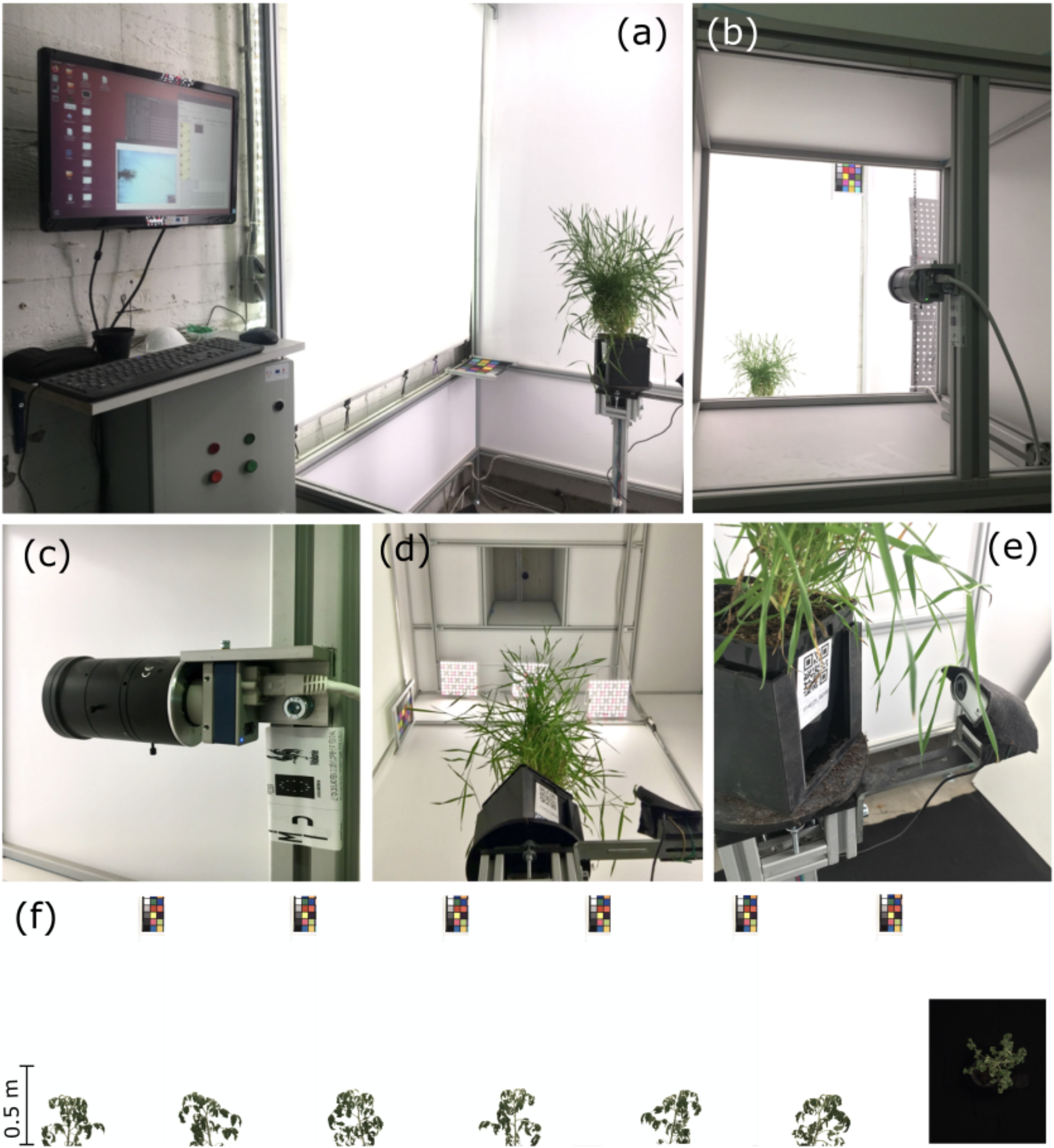
Views of the imaging cabinet including (a) the acquisition interface, (b,c) the side-view camera, (d) the rotating platform and the top-view camera, and (e) the QR-code reading webcam (bottom right). Typical images (f) obtained with the system including 6 side-views obtained by rotating the plant (white background) and one top-view (black background). A reference color chart was used to calibrate the images (upper right corner in the side-view images, not shown for the top-view image).

The cameras and the stepper-motor were controlled through a dedicated software written in Python and running on a Linux computer in order to synchronize plant identification, rotation, and image acquisition. The adjustment of basic camera settings (e.g. shutter speed, gain, output format) used libraries from OpenCV (https://pypi.org/project/opencv-python/) and Aravis (https://github.com/AravisProject/aravis) while rotation functionalities (i.e. speed, number and time of acquisitions after QR-code detection) was programmed by us. Typically, six side-view images and one top-view image were acquired during a 180° rotation in 4 seconds (45° per sec.). The plants were manually loaded on the rotating platform through a sliding door, which was closed before imaging. After rotation was initiated, the imaging cycle started when the QR-code identifier was read by the webcam, and each image acquired by the Genie-Nano cameras was saved under a unique ID. After each imaging, a preview of the images allowed a quick visual check and, in case of a problem, a second imaging cycle was performed. The complete imaging cycle was about 10 seconds per plant, allowing to image the complete set of 348 pots included in this experiment in less than half a day.

The image acquisition software saved each image under a unique ID. An automated processing script was developed using the macro language of the ImageJ open source image processing package (Fiji distribution) (46). This script was used to extract plant phenotypic descriptors from each image through a fully automated process, including the following steps: i) read the raw image in bayer format, ii) get the metadata (e.g. date, plant ID, camera view, frame number), iii) perform white balance and spatial calibration based on a reference color chart, iv) segment the plant from background using grey-scale or color thresholding, v) measure plant dimensions and shape factors, vi) extract color components in either RGB or HSB color space, vii) export raw data in text format (.csv).

A more detailed description of the image processing is available as supplemental material (Table S1).

The output data included three types of phenotypic descriptors: i) simple dimensions (e.g. Height, Width, Projected Area, Fitted Ellipse), ii) shape factors derived from dimensions (e.g. Roundness, Solidity, Circularity), and iii) color density and variation values (Red, Green, Blue, Hue, Saturation, Brightness, and their respective standard deviations). For each species and timepoints, these parameters were saved into separate text files that were ultimately combined into one large dataset.

### 2.6 Data processing and analysis

R version 3.6.1 for macOSX (available at https://cran.r-project.org/bin/macosx/) running under Rstudio version 1.0.136 (Rstudio, Boston, MA, USA) was used to: i) compute additional shape factors as ratios from existing measurements, such as Voxel, Compactness, Anisotropy; ii) compute color indices such as Green Leaf Index (GLI), and Triangular Greenness Index (TGI); iii) generate a chlorophyll content prediction based on RGB values; iv) generate scatter plots to visually check for abnormal measurements (e.g. corrupted images) before further statistical use; v) aggregate the multiple camera measurements per plant (e.g. the side view camera generated 6 images from which the mean, maximum, minimum, and median values were computed); vi) merge imaging data with plant metadata (species, spatial location, light intensity and quality at plant location, measured chlorophyll content); vii) evaluate species discrimination based on Principal Component Analysis (PCA); viii) apply linear regression to quantify the effect of the Red:Blue gradient on each phenotypic parameter at successive timepoints.

A detailed description of the phenotypic descriptors and how they were calculated is available as supplemental material (Table S2).

For side-view data, since there were 6 different images per plant, we had to choose whether to use the mean, the median, or another statistics. After some testing we decided to use the mean of the 6 images, except for the parameters Height and Width for which we used the maximum values.

At each timepoint, and for every measured parameter, linear regression with the Red:Blue ratio was used to generate correlation coefficients such as Pearson R, p-value, slope, and intercept. We decided to use the simplest possible linear model (y = a*x + b) to estimate the “effect size” of the gradient as the percentage of the difference across the Red:Blue gradient (Effect size (%) = (value at max PFD_Red_:PFD_Blue_ - value at min PFD_Red_:PFD_Blue_) / value at min PFDRed:PFD_Blue_ * 100). The slope and the intercept were used to compute value estimates for each phenotypic parameter at the minimal and the maximal Red:Blue ratios (value estimate = slope x Log10(PFD_Red_:PFD_Blue_) + intercept).

### 2.7 Chlorophyll content measurements

The leaf chlorophyll content was estimated with an Apogee MC-100 (Apogee Instruments, Logan, UT, USA). Built-in calibration models were used for tomato and rice, whereas the built-in generic model (https://www.apogeeinstruments.com/content/MC-100-manual.pdf) was used for the other species. At least 6 measurements were made on minimum 3 different mature leaves per pot. The area of measurement was located on the most horizontal part of the limb, which was the most exposed to light, and the measurements were distributed across its width. The measurements were averaged per pot.

## 3. Results

### 3.1. Phenotypic discrimination of species based on image data

Seven different species were grown under a gradient of Red:Blue light. Four species were dicots, of which one rosette (*A. thaliana*), one caulescent (*S. lycopersicum*) and two bushes (*E. peplus* and *O. basilicum*). The three other species were monocots, all tillering rosettes with erect leaves (*O. sativa, B. distachyon, S. viridis*). Changes in growth, morphology and color were recorded, based on plant imaging every 3-4 days. Three types of phenotypic descriptors were collected: i) dimension, ii) shape factors, iii) color indices. For each type of descriptor, different proxies were extracted from images captured from both top- and side-view cameras (see the Materials & Methods section for more details).

In order to evaluate the performance of our phenotyping setup and to select the most discriminant plant features, PCA analyses were performed using either dimension, shape factors, or color indices only, or all parameters together (Figure 4). Imaging data collected over 3 timepoints between 21 and 29 days after transfer under Red:Blue gradient were used to generate the PCA plots.

**Figure 4.**
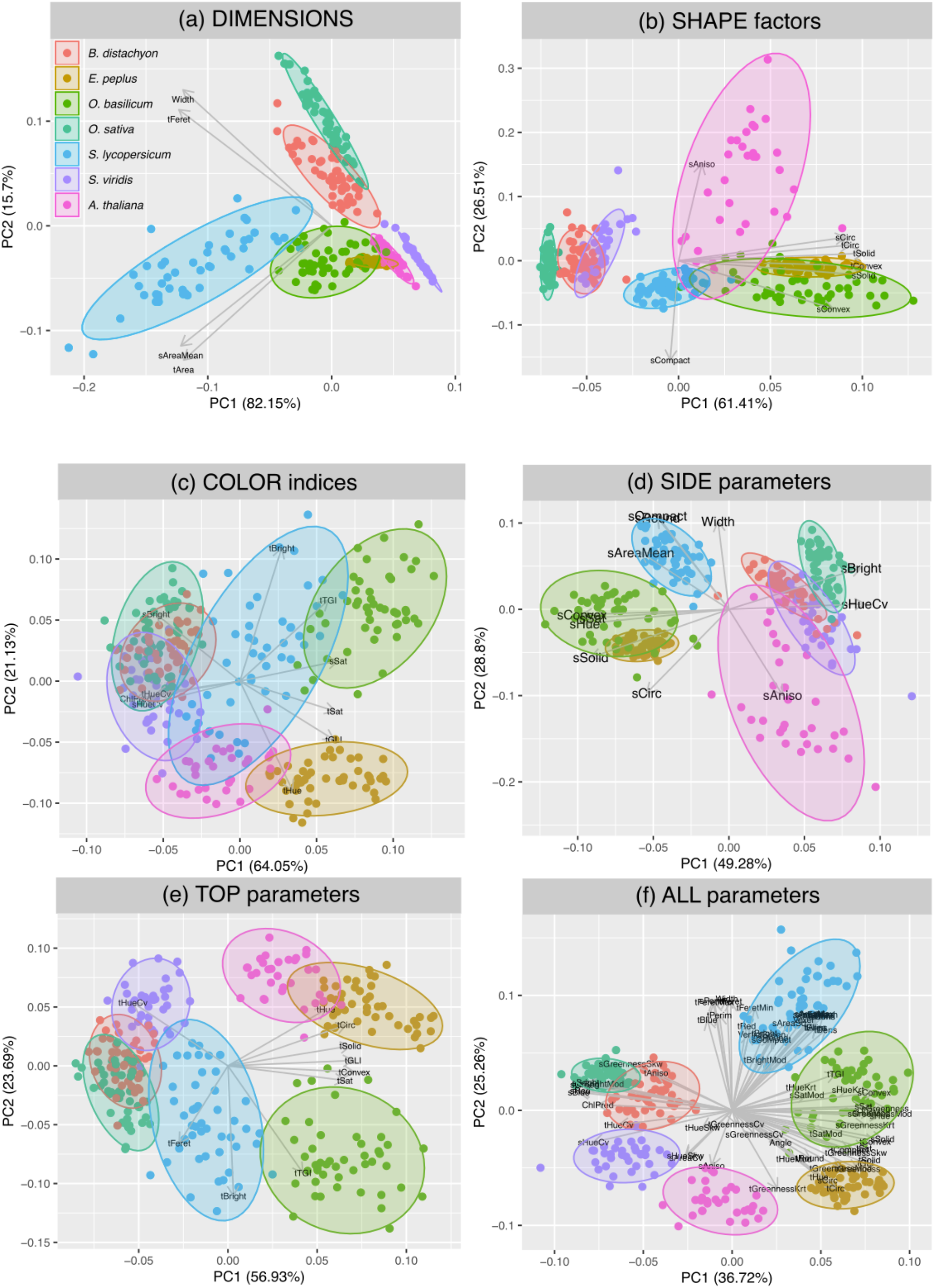
Principal Components Analysis discrimination of seven species based on various selections of measured parameters. Species color codes are shown in panel (a).

The different species can be discriminated based on simple dimension features (Figure 4a), except *E. peplus* and *O. basilicum* that overlap completely. Shape factors alone neither separate clearly *B. distachyon* from *S. viridis* nor, again, *E. peplus* from *O. basilicum* (Figure 4b). Color indices alone do discriminate rather well the 3 dicots but not so much the 3 monocots (Figure 4c). When all 3 types of parameters are combined, a much better separation can be achieved with monocots and dicots clearly pulled apart in opposite sectors of the PCA plot (Figure 4f). Finally, Figures 4d-e show that using only top-view or only side-view data yields different separations, e.g. *E. peplus* and *O. basilicum* separate well with top-view traits but not with side-view traits, while the opposite stands true for *O. sativa* and *B. distachyon*. Therefore, both side- and top-views are needed to achieve the best discrimination. Note also that *A. thaliana* data show greater variation than the other species for shape factors (Figure 4b) and side-view traits (Figure 4d). This can be explained by bolting occurring in the time-course of the experiment and affecting shape factors such as Anisometry and Circularity, which are sensitive to changes in overall symmetry and elongation. For this reason, side-view data for *A. thaliana* were not used in the following analyses.

### 3.2. Gradient effects on plant phenotypes

The Red:Blue ratio (PFD_Red_[600-700nm] over PFD_Blue_[400-500nm]) was calculated for individual plants based on the spectral light measurements performed at each plant position at the beginning of the gradient treatment. For plants growing under white light, it was calculated from an average of several measurements. The Red:Blue ratio affected the three types of phenotypic descriptors that we measured - dimension, shape and color - but at various extent and sometimes in opposite directions in the different species. Figure 5 shows two examples of eye-perceptible effects. *S. lycopersicum* plants (photographed 20 days after start of the gradient treatment) were clearly taller and wider as the Red:Blue ratio increased, due to stem and petiole elongation. These phenotypes were detectable with dimension and shape proxies, as described below, but also with color changes due to more stem and petiole parts being exposed. By contrast, no clear effect on rosette size was detected in *A. thaliana* plants (photographed 12 days after start of the gradient treatment) but its shape was changed due to curling of the leaf margins under exposure to Red light (Figure 5, panels (e) and (f)). No color change was perceptible by eye.

**Figure 5.**
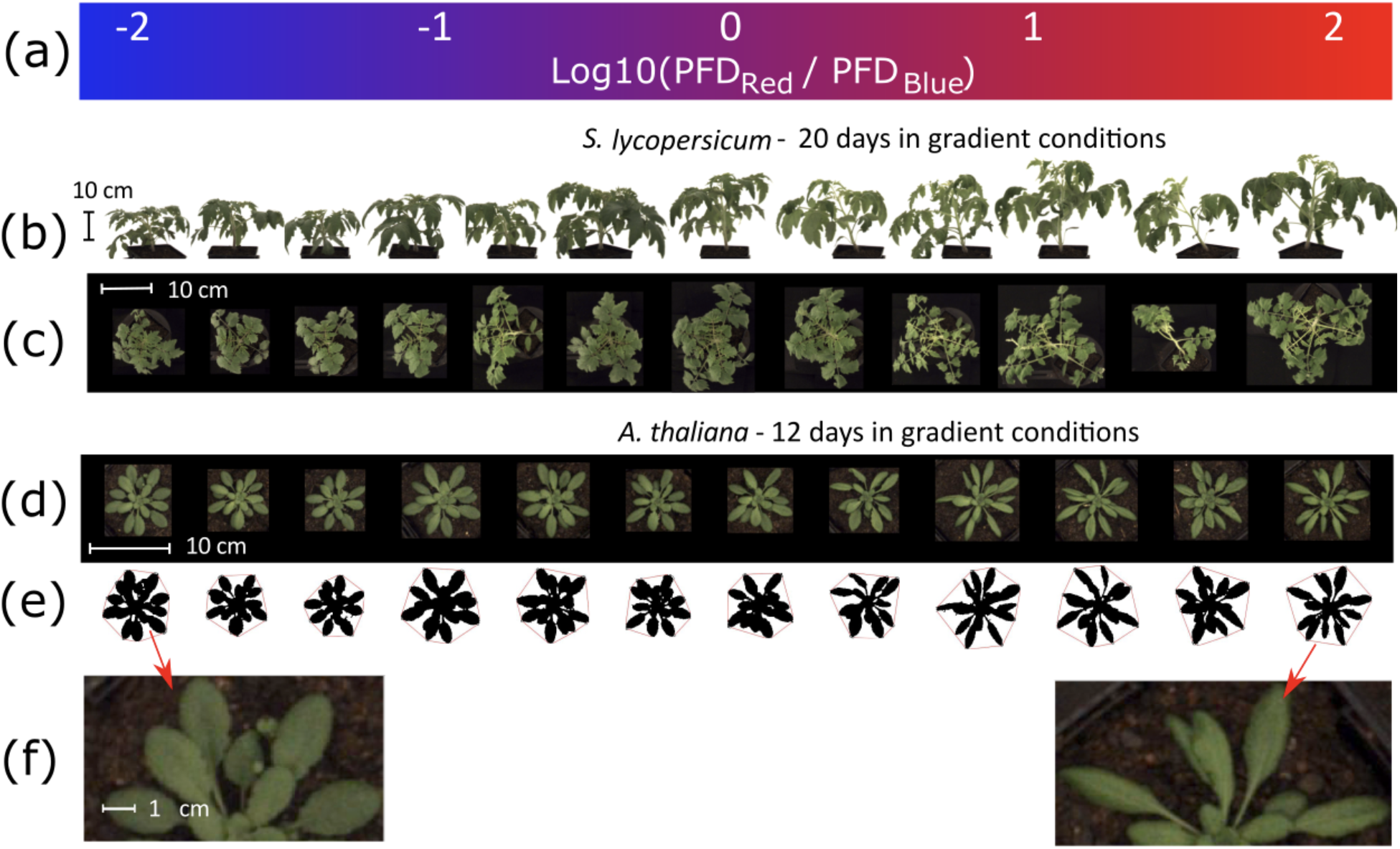
Example of plant phenotypes under the Red:Blue gradient. (a) light gradient, (b) side-view images of a row of *S. lycopersicum* plants, (c) top-view images of the same *S. lycopersicum* plants, (d) top-view images of *A. thaliana* plants, without thresholding, (e) same images of *A. thaliana* plants, after thresholding (red line is the convex hull of the object), (f) enlarged images of the *A. thaliana* plants on each extreme side of the gradient.

The effects of the light gradient were quantified by plotting phenotypic measurements versus the logarithmic value of the Red:Blue PFD ratio, calculating linear regressions, and computing correlation coefficients (R^2^, p-value, slope, and intercept).

Figure 6 shows examples of such linear regressions for a few parameters that responded strongly to the Red:Blue ratio. These examples show that the responses may be different between species, even sometimes opposite. For example, plant height increased with higher Red:Blue ratio in most species, except in *O. basilicum* and *S. viridis*. Projected leaf area measured with top-view imaging was also correlated with increased Red:Blue ratios in two species, *B. distachyon* and *E. peplus*, while the other species showed little or no significant change. Circularity, a shape factor that quantifies Area:Perimeter variations, was decreased under higher Red:Blue in at least two species, *E. peplus* and *S. lycopersicum*, but increased in *O. basilicum*. In *E. peplus* and *S. lycopersicum*, this was likely due to stem and petiole elongation, which reduced leaf overlaps and thus created gaps in the canopy, therefore reducing Circularity that is sensitive to the number and size of concavities in the contour of the measured object. On the contrary, in *O. basilicum*, such gaps were present but decreased as leaves grew.

**Figure 6.**
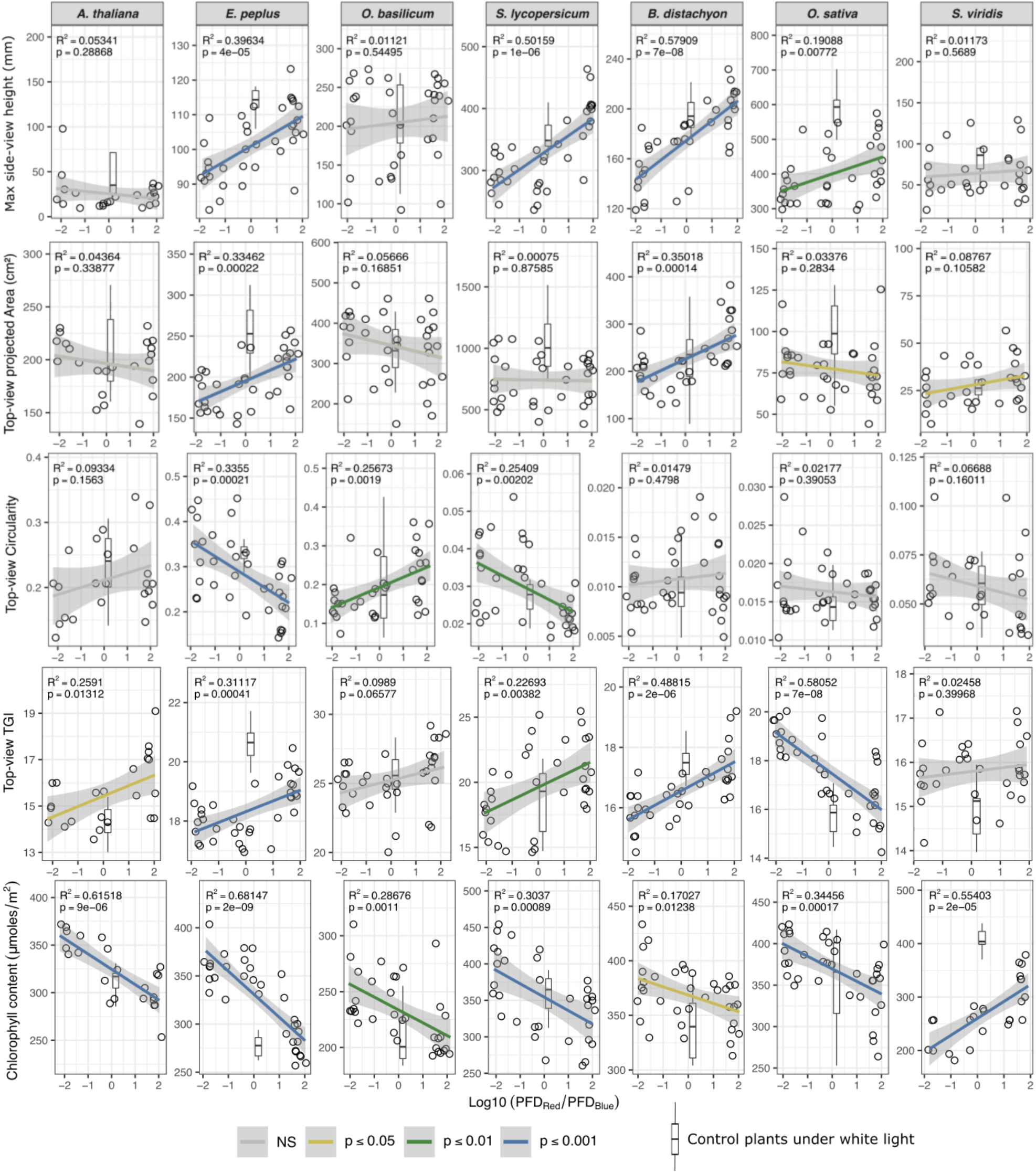
Examples of linear regressions between the phenotypic traits measured 29 days after the start of the gradient treatment and the Red:Blue ratio. Each point is an individual pot with one plant (*A. thaliana, S. lycopersicum, B. distachyon, O. sativa, S. viridis*) or one bush of several plants (*O. basilicum, E. peplus*) (see Materials and Methods). Greyed areas are 95% confidence intervals. Boxplots represent minimum and maximum values (whiskers), median (horizontal line), first and third quartiles (box).

The triangular greenness index (TGI) increased with higher Red:Blue ratios in all species except in *O. sativa*. TGI is a calculation based on Green reflectance relative to Red- and Blue- and has been reported to be negatively correlated to chlorophyll content (47). This negative correlation is explained by the fact that when chlorophyll concentration increases, leaves appear darker and hence reflectance values decrease. The effect of the gradient on TGI was coherent with leaf chlorophyll content measured with a chlorophyll meter in 5 out of the 7 species (Figure 6). The two exceptions were *O. sativa*, where both TGI and chlorophyll content decreased with the Red:Blue ratio, and *S. viridis*, where the chlorophyll content increased with the Red:Blue ratio while the TGI showed little variation. The decoupling of TGI and chlorophyll in these two species might be due to color changes involving other pigments or could be explained by a shape effect affecting leaf reflectance. Another surprising observation in Figure 6 is the effect of the Red:Blue gradient on the chlorophyll content of *S. viridis*, which is completely opposite to what was observed in the other species. Since *S. viridis* is the only C4 species in the experiment, it is tempting to suggest that C4 and C3 plants might differ in their response to the light spectrum. Finally, it can be seen in Figure 6 that the plants growing under standard white light sometimes differed phenotypically from plants grown under LEDs at the same Red:Blue ratio. For example, in *E. peplus*, all measurements shown in Figure 6 under white light are outside the confidence interval of the Red:Blue gradient. These observations clearly indicate that plant phenotype under the white light conditions was affected by other factors than the Red:Blue ratio, but to various extent in different species.

As described in section 2.6, we summarized the bulk of phenotypic data by performing, at each time point and for each species, regression analyses for every phenotypic descriptor versus the Red:Blue ratio and extracted the coefficients R^2^, p-value, slope, and intercept. Then, we computed phenotypic values at the lowest and highest Red:Blue ratios using the slope and intercept of the regressions to evaluate the size of the gradient effect. In Figure 7, effect sizes are presented as the relative difference (% effect) between the extreme sides of the gradient after 29 days. In this figure, we selected 20 parameters, out of >30 measured, for which a highly significant correlation (p<0.01) with Red:Blue ratio was found in at least one species. Note that the sign of the effect size is arbitrary since it is determined by the direction of the Red:Blue gradient. A figure presenting the corresponding R^2^ values is also available as supplemental material (Figure S2).

**Figure 7.**
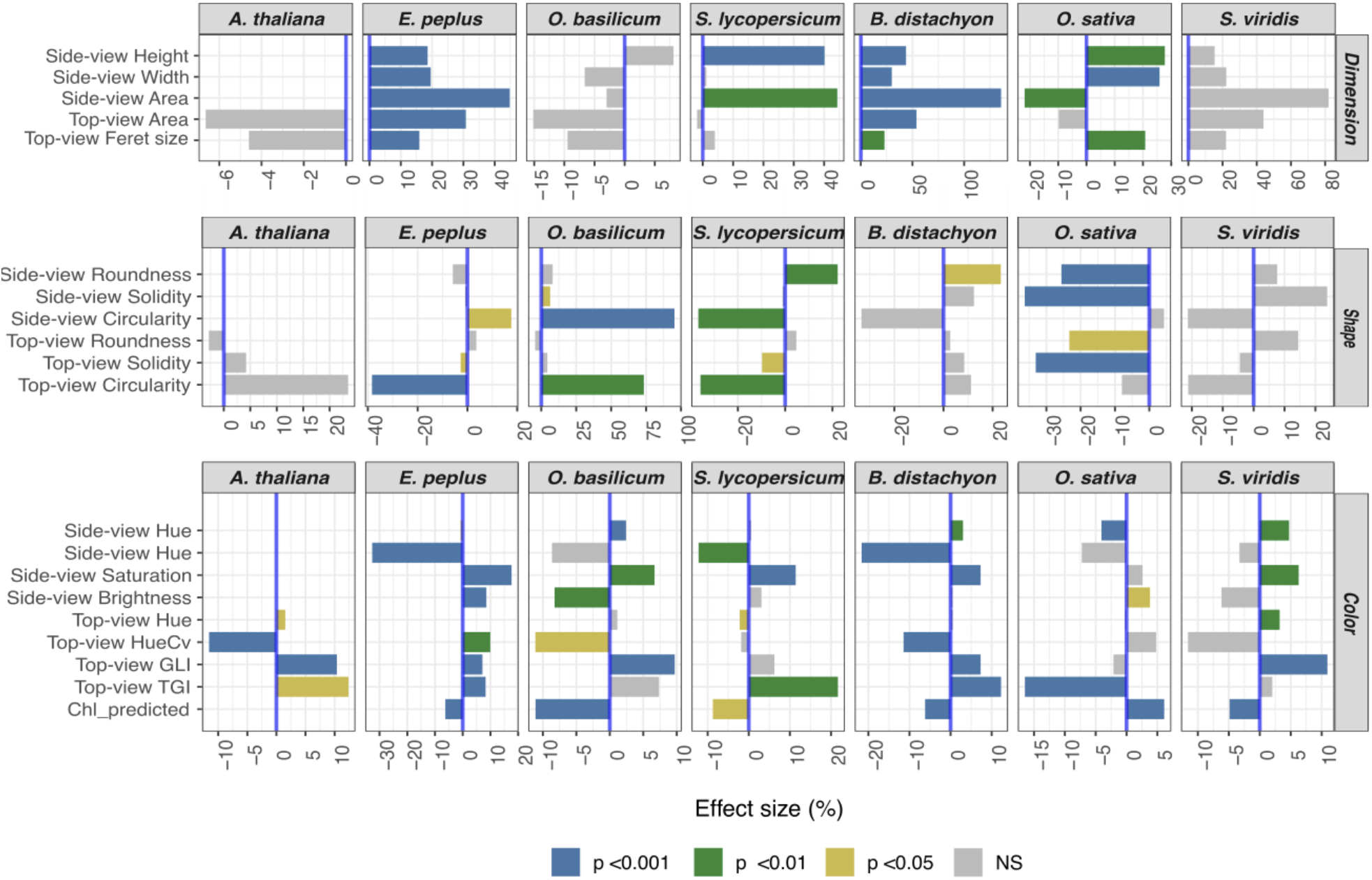
Red:Blue gradient effect size (%) measured 29 days after the start of the gradient. Effect size was computed as explained in Materials and Methods. Only 20 parameters are shown, for which a highly significant correlation (p<0.01) with Red:Blue ratio was found in at least one species. The significance categories are based on the p-value of the computed R^2^.

In terms of plant size, a correlation between plant dimensions (Area, Height, Width) and higher Red:Blue ratios was clearly seen in all species, except in *O. basilicum*. In *A. thaliana* and *S. viridis*, the changes did not reach significance, possibly due to higher variability or, in the case of *A. thaliana*, due to maximum size of the rosette being reached in all conditions.

The variations in the measured shape descriptors were more complex, with species-specific patterns. Here we focused on 3 parameters that respond to different “behaviors”: i) Roundness as defined in ImageJ is the ratio between the fitted ellipse’s minor and major axes and decreases with elongation, ii) Circularity is the ratio of the object’s area to the area of a circle having the same perimeter and decreases as concavities increase, but is relatively insensitive to contour roughness, iii) Solidity is the ratio of the object’s area to the convex-hull area and decreases with rough contours and holes.

In *A. thaliana*, *S. viridis*, and *B. distachyon*, no or very few significant changes could be recorded amongst the 3 selected shape descriptors, while in the other four species we observed clear effects, as illustrated in Figure 8. The main effects of the Red:Blue ratio on plant shapes were: i) a strong increase of Circularity in *O. basilicum* due to the stems of the bush being more tight together, ii) a decrease of Circularity in *S. lycopersicum*, for both side and top view images, likely caused by stem and petiole elongation, and also in *E. peplus*, but only in the side-view images, iii) decreased Roundness and Solidity with more erect leaves in *O. sativa*. Note that in this species, Circularity values are very low because of the very narrow leaves, making this parameter less reliable.

**Figure 8.**
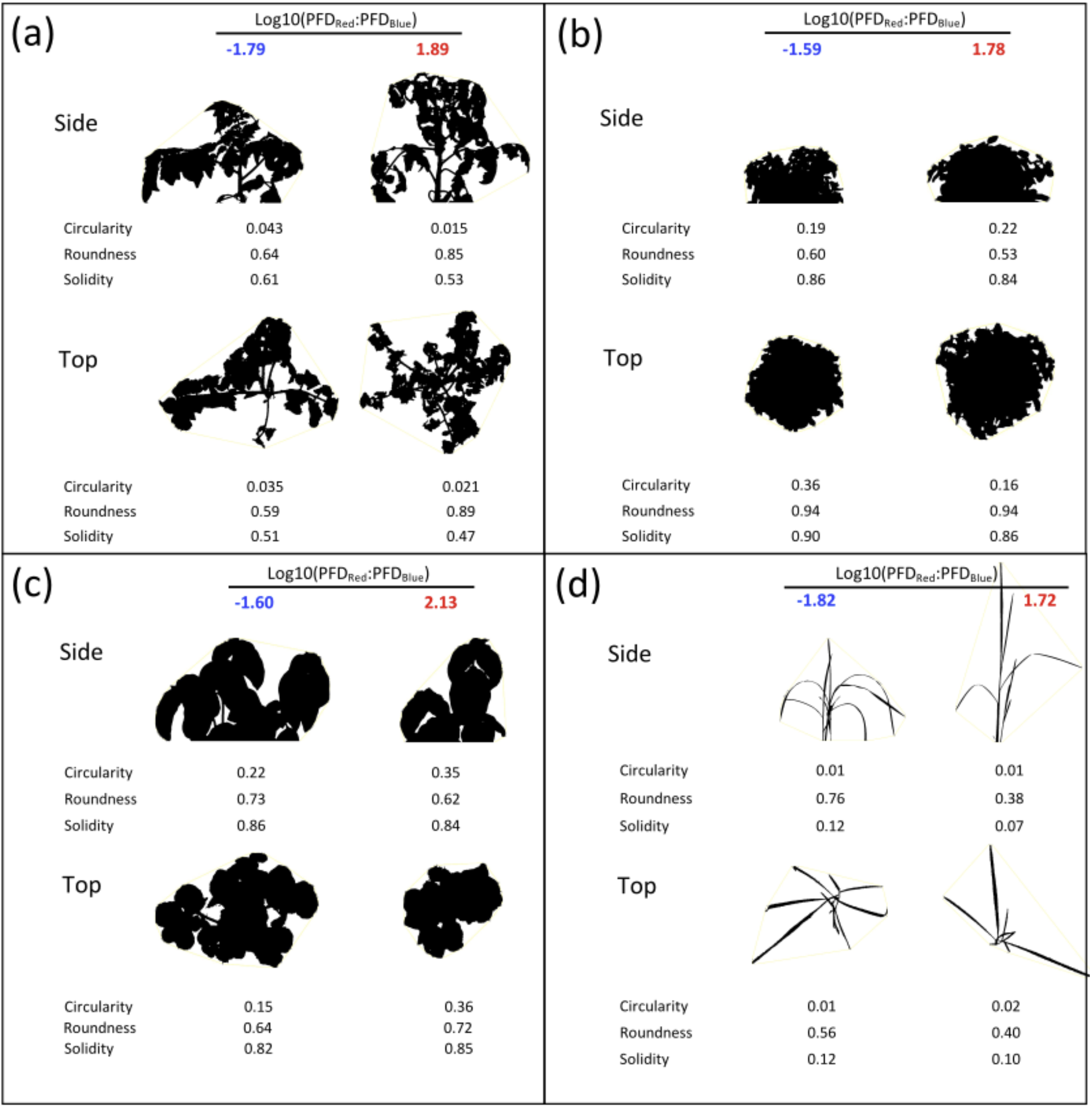
Individual plant silhouettes illustrating the effects of the Red:Blue gradient on plant shape factors in four species: (a) *S. lycopersicum*, (b) *E. peplus*, (c) *O. basilicum*, (d) *O. sativa*.

Finally, changes in the color indices seemed more consistent across the panel of species: i) the two greenness indices, TGI and GLI, increased with higher Red:Blue ratios, with the notable exception of *O. sativa*; ii) the coefficient of variation for Hue recorded from the side-view picture decreased with the Red:Blue ratio, though not always significantly, indicating a more uniform tone of color under higher Red:Blue ratios; iii) the Saturation in the side-view images increased, which could be the sign of denser pigmentation under higher Red:Blue. This increased pigmentation would then be due to other pigments than chlorophyll since leaf chlorophyll measured with the transmission probe followed an opposite trend (Figure 6).

Finally, we plotted the evolution of the slopes of the linear regressions that were calculated between phenotypic traits and the Red:Blue ratio at different time points during and after the gradient treatment (Figure 9). The purpose is to visualize how the effects of the Red:Blue gradient evolved in the time course of the experiment and whether they were maintained or not after returning the plants to white light.

**Figure 9.**
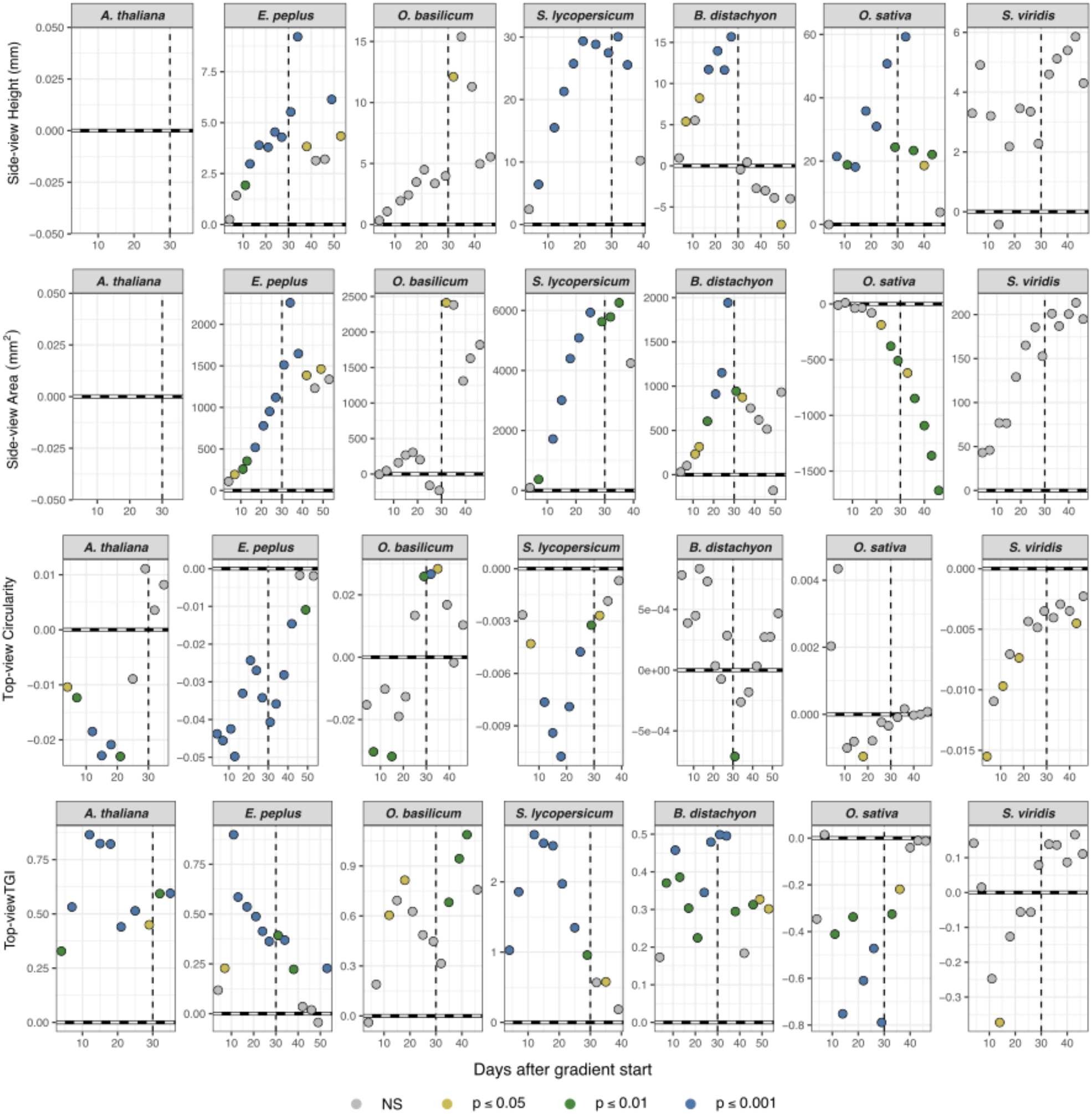
Evolution with time of the phenotypic response to the Red:Blue gradient. Values shown are the slopes of the regressions between phenotypic traits and the Red:Blue ratio, calculated at each time point. Horizontal solid line: slope = 0 (no effect of the gradient). Vertical dotted line: end of the Red:Blue gradient treatment, return to white light. The significance categories are based on the p-value of the computed R^2^.

Figure 9 shows that the effects of the Red:Blue gradient on plant dimension descriptors (side-view Height, Area, and top-view Area) were measurable soon after start of the treatment and increased for its duration. For example, the side-view Height increased more and more with the Red:Blue ratio and with the time spent under the gradient. This phenotypic change was reversible under white light in *B. distachyon, E. peplus, O. basilicum*, and *O. sativa*, indicating that the differences in height were due, at least partly, to changes in plant stature, i.e. changes in shoot and leaves inclination. In *B. distachyon* and *E. peplus*, the side and top Area increased with the same trends than side-view Height, indicating a reversible opening of the plant bush under higher Red:Blue ratios. The opposite was observed in *O. sativa*, with the slope of the correlation for side and top Areas decreasing gradually during the gradient treatment and even after retransfer to white light. This behavior can be explained by a more erect position of the leaves under higher Red:Blue ratios, which is consistent with the observation that rice was the only species with decreased Roundness (more elongated shape) at high Red:Blue ratio (see Figures 7 and 8).

In *S. lycopersicum* and *S. viridis*, the increases in side-view Height, side-view Area, and top-view Area with the Red:Blue ratio were maintained after the treatment, reflecting irreversible changes in plant growth with light quality. Internode elongation was indeed observed in both species.

Shape and color factors showed abrupt variations after the start of the Red:Blue treatment in all species. In some cases, the synchrony of the changes clearly suggested a correlation between traits. For example in *S. lycopersicum*, changes in top-view Circularity and top-view TGI appeared almost perfectly synchronized, indicating a high correlation - in this case, negative - between these parameters. This observation suggests that plant shapes may influence color indices through a change of reflectance. However, color indices such as TGI may still be indicative of pigment composition as we observed an inverse correlation with chlorophyll content in 5 out of 7 species (see Figure S3), with only *O. sativa* and *S. viridis* showing slightly positive or no correlation, respectively.

## 4. Discussion

The combination of new LED technologies with high-throughput phenotyping pipelines provides unprecedented perspectives for research and agricultural applications. The purpose of our research was to explore the capabilities and possible applications of combining smart LED devices with image-based phenotyping for the characterization of model plants and crops. Therefore, we selected seven species for simultaneous experimentation. While most studies focus on monochromatic LEDs, our approach showed that growing plants under a color gradient provides reliable data with additional insights, in the form of correlative trends, which are key to control plant responses and eventually ameliorate desired characters. Indeed, spectrally variable LED lighting sources allow flexible set up, including the use of distinct light recipes that can be tested simultaneously within the same cabinet, with light quality as the only explanatory variable to the observed phenotypes.

To evaluate the impact of a Red:Blue light gradient across time, we acquired several images for each individual plant at regular intervals. These images were analyzed using an automated analysis software that retrieved multiple measurements, including plant dimensions, shape factors, and color indices. These measurements or phenotypic descriptors, are different from those obtained using ruler-based visual methods. Although they are not always easy to apprehend, they have the key advantage to be highly precise, thus allowing the detection of subtle changes that would not be noticeable by an operator, such as shifts in shape or color that are not visually obvious. In order to make sense of the very large dataset that we generated, we used a correlative approach and calculated linear regressions between many dependent variables —the phenotypic descriptors— and one explanatory variable —the Red:Blue ratio. The resulting regression coefficients allowed us to grasp the response profiles of the different species exposed to the color gradient.

### 4.1. General effects of the light gradient

A key question of this study was whether we would identify common trends within the response of different species to the light gradient. To investigate this question, we quantified three categories of phenotypic descriptors in the selected seven plant species: dimension and shape indices are relevant to quantify variations in the plant stature, while color indices indicate possible variations in plant pigments. It is important to note that, although color indices (e.g. TGI) can help detect differences that are not visually obvious, they can be ambiguous and do not always accurately reflect actual changes in pigment contents, so that further species-specific calibrations are required before any practical application. The time-course analysis of dimension, shape, and color indices during and after the light treatment revealed that some effects of the gradient were reversible, either within days or more slowly, while others were irreversible. We found responses to the light gradient for most species, but the amplitude and direction of these changes were remarkably species-dependent (Figure 10). Consequently, general phenotypes cannot be predicted without experimental work, highlighting the need to analyze each species separately.

**Figure 10.**
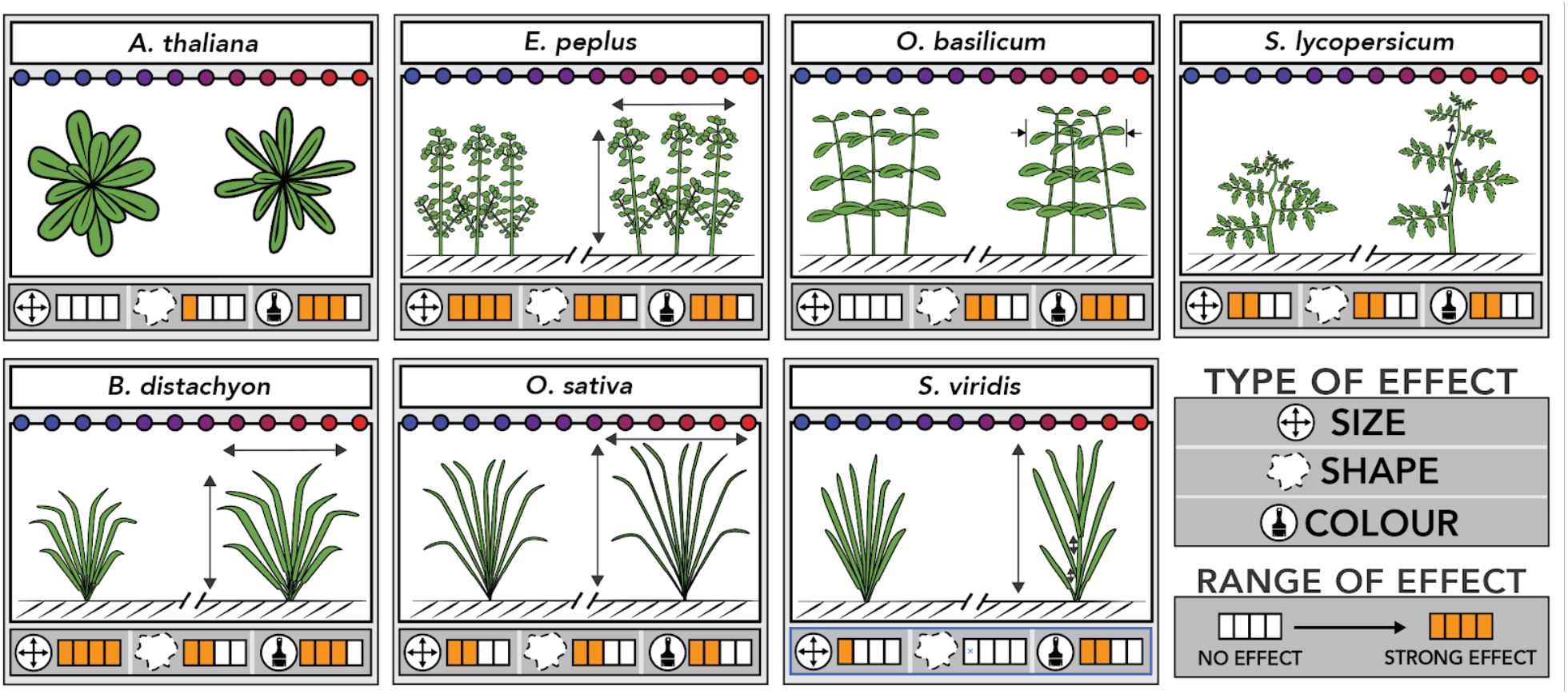
Schematic representation of the phenotypic variations caused by a Red:Blue light gradient in seven plant species. Effects observed 29 days after the start of the light gradient.

### 4.2. Species-specific behaviors

In *A. thaliana*, we did not observe any effect of the light gradient on the size of the rosette. However, we detected that higher Red:Blue ratios caused an increase in the top-view Circularity parameter. This change is likely caused by an increase in leaf curling, a known red light-induced phenotypic response. Indeed, Inoue et al. reported that, upon exposure to red light, newly initiated leaves were curled and slanted downward, a phenotype that could be reversed by the addition of blue light (14). A similar phenotype, known as the “red-light syndrome”, has been reported in other species, including crops (29, 48).

*E. peplus* is an annual medicinal eudicot whose sap, which is toxic to rapidly replicating human tissue, has long been used as a traditional remedy for common skin lesions and, more recently, for pre-cancerous pathologies. To the best of our knowledge, this is the first study involving the indoor cultivation of this species and our observations suggest a potential mean to optimize its biomass production. Indeed, *E. peplus* was the species that responded the most homogeneously in terms of plant dimensions, as all size-related descriptors were increased under higher Red:Blue ratios. This phenotype was the consequence of an increased growth of the bush in all directions. Stem elongation loosened the compact foliage, thus decreasing side-view Circularity under higher Red:Blue ratios.

In *O. basilicum*, we did not identify any significant effect of the light gradient on size descriptors, but we did observe an increase in the Circularity shape factor, which suggested a compaction of the bush under higher Red:Blue ratios. Previous reports, in which the response of *O. basilicum* to environmental factors was assessed using biomass measurements, showed conflicting results: blue light was reported to either improve (49, 50) or to reduce shoot biomass by limiting stem elongation and leaf expansion (51), depending on the growing setup. Our results, however, show that the shape of *O. basilicum* bush can be manipulated by light, which can be a valuable tool to meet market specifications.

In *S. lycopersicum*, we observed a strong increase in shoot height under higher Red:Blue ratios. This phenotype, which is a consequence of higher internode elongation, is consistent with previous studies showing that blue wavelengths inhibit stem elongation in phylogenetically distant eudicot species such as lettuce, soybean, and tomato (52–54). We also found differences in color proxies (TGI, chlorophyll measurements) along the color gradient, which are consistent with a previous report showing that blue light exposure increases chlorophyll content in tomato leaves (54).

Among the three monocots that we studied, *B. distachyon*, a species increasingly used as a model plant to study developmental processes in Pooids, is the organism whose dimensions were the most affected by the light gradient. We observed an increase in height and diameter upon increasing Red:Blue ratio. One possible explanation would be a reduction of leaf length by blue light, as already reported for wheat (55). Another possible explanation would be a stimulation of tillering upon red light exposure, but closer observations are required to test that hypothesis.

Interestingly, in *O. sativa*, we observed that increased Red:Blue ratios alter the plant shape by enhancing the erectness of leaves and causing plant tightening, as reflected by changes in the Roundness and Solidity phenotypic descriptors. These modifications of the plant stature could explain that the color indices based on leaf reflectance (e.g. TGI) were not good proxies of chlorophyll content in this species. Interestingly, erect leaves were previously shown to improve photosynthesis and yield in rice by reducing leaf shading in dense plantations (56). This phenotype is regulated by environmental and hormonal factors, among which brassinosteroids exert a prominent role. The effects of light quality observed here could act upstream of these hormones, as suggested in earlier reports (57).

In *S. viridis*, a Poaceae used as a model to study C4 photosynthesis, we observed unexpected effects of the light gradient on the chlorophyll content. Indeed, whereas the chlorophyll content decreased with higher Red:Blue ratios in all other species, it increased in *S. viridis*, but remained lower than under white light, again in opposition to other species. It is tempting to speculate that this peculiar behavior of *S. viridis* is linked to its C4 metabolism, but there are actually not many other reports that we are aware of to confirm this idea. In one report, though, it was shown that, in maize, blue light represses the accumulation of chlorophylls, compared to red light (58), which seems consistent with our observations. On a different level, it is noteworthy that this effect on chlorophyll in *S. viridis* was not revealed by color indices such as TGI, illustrating the limitations of non-destructive color proxies. One explanation is that, unlike in the other species, *S. viridis* plants started flowering during the gradient treatment, and TGI may have been biased by the presence of paler green panicles, independently of the variations in leaf chlorophyll content.

In conclusion, the effects of the Red:Blue gradient are strongly species-dependent and do not allow generalization. It would be interesting, however, to broaden the analysis to more plant species to test whether functional groups showing similar behavior can be identified.

### 4.3 Future Improvements and Perspectives

The pipeline presented here proved to be effective to screen the effects of a light gradient on the phenotype of multiple plant species. Some technical components could, however, be improved. For example, our low-cost in-house imaging station requires the manual transport of plants, and hence an automated conveyor system would reduce operating time by an estimated two fold at least. Alternatively, an in-chamber top view imaging device could be used although with its own caveats, additional analysis challenges, and limitations. In particular, it is not well suited to phenotyping individual plants within a canopy, which is a major statistical drawback.

In order to validate our pipeline, we have chosen a simple light mixture of red and blue lights, which has been the focus of many publications in the horticultural domain. However, many types of gradients can be tested, including linear gradients involving other wavelengths (Red:Far-Red, Red/Blue:Green, UV-A:UV-B) and bi-dimensional gradients, which would help explore a larger number of conditions in a single experiment. The gradient approach could also be used to determine the optimum of light mixture required for a given trait, or to acquire the data necessary for modeling plant responses to the light quality.

Image analysis was performed using the popular, and free to use, generalist package, ImageJ. The same measurements could also be accomplished with many other available softwares, some of which offer more specialized functionalities for plant phenotyping (see https://www.quantitative-plant.org/software for an overview of available plant phenotyping applications). Nevertheless, our process turned out to provide exploitable proxies for plant dimension and shape, although color indices were not always correlated with differences in chlorophyll contents. Indeed, the color indices may be, at least partially, sensitive to differences in the reflectance caused by distinct plant shapes and leaf orientations. Similar issues were previously reported in studies on spectral imaging and the solution requires capturing leaf orientations and subsequently modeling light reflectance (59).

The accuracy, relevance, and depth of phenotyping could be improved by using new imaging technologies such as spectral, tridimensional, thermal, or fluorescence imaging, depending on the desired application. Additional calibration steps based on conventional biometric measurements of plant biomass remain highly recommended to ascertain the significance of imaging-based phenotypic descriptors. Another attractive approach is to use machine learning to facilitate the interpretation of the complex set of parameters generated by imaging, especially when it comes to phenotypic descriptors such as shape factors and color indices which are more difficult to grasp. For example, classification techniques would allow to categorize plants according to pre-defined criteria, and provide the user with a more holistic understanding of the plant phenotype.

### 4.4 Conclusions

To conclude, the setup described here can be improved and upscaled in many of its aspects to meet a variety of research needs. Still, the unique combination of light gradients with in-depth phenotyping obviously provides new perspectives to address fundamental plant biology questions as well as help improve applications in screening, breeding, modeling, or functional genomics. In particular, this approach provides innovative tools for the development of new varieties that are better suited for indoor light conditions.

## General

Early access to the Lumiatec LED lighting systems was possible with the collaboration of GDTech and Araponics R&D teams, especially George Ferdinand, Michaël Menu, Julien Reuland, and Dylan Dohogne. The authors are also grateful to Sébastien Steyaert and Gabriel Berger, for their technical assistance for plant cultivation and imaging, and to Profs Frédéric Lebeau, and Guillaume Lobet for fruitful discussions and comments on the manuscript.

## Author contributions

PL, AF, FB, SHF, PT, and CP designed the experiment and wrote the paper. AF and PT setup the Lumiatec luminaries. PL created the imaging cabinet; AF and PT developed the image acquisition software and interface. PL designed the image analysis script and processed the raw data.

## Data Availability

Data, including measurements database, image analysis script, R script, and link to raw images, are available at zenodo.org (https://doi.org/10.5281/zenodo.4071811).

## Supplementary Materials

**Table S1.** Steps in the image processing to generate plant morphology and color measurements.

| Nr | Description | Comment |
| --- | --- | --- |
| **Steps in ImageJ** |  |  |
| 1 | Read image file, get plant name, camera view, and frame number | "_0_" = side view "_1_" = top view |
| 2 | Find blue square in the color target and extract x,y coordinates | To convert to HSB color model, threshold light blue objects and record x,y coordinates |
| 3 | White balance using grey values on the reference target card | Adapted from P. Mascalchi (https://github.com/pmascalchi/ImageJ_Auto-white-balance-correction) |
| 4 | Set the ROI (region of interest) | To remove borders and reference card |
| 5 | Create a HSB (hue, saturation, intensity) image | The HSB image is used later for color measurements |
| 6 | Separate RGB channels into 3 grey-level images | For both thresholding and computing greenness indices |
| 7 | Side-view images only:<br>Threshold on the B (blue) channel | To segment the plant from the background before measurements |
| 8 | Top-view images only:<br>Color threshold in HSB (hue, saturation, brightness) color space | Color thresholding is much slower than single channel thresholding, but is necessary when the background is not uniform as is the case with top view images |
| 9 | Eliminate small noise blobs based on size threshold | To eliminate any small background artifacts |
| 10 | Erode irregularities around the segmented object shape | To increase precision of contour measurements |
| 11 | Create a selection for morphological measurements | This is a binary “mask” of the plant |
| 12 | Save cropped color image for visual check | For rapid post-processing visual checks the smallest region enclosing the plant is saved as a separate color image |
| 13 | Measure plant dimensions on the segmented shape | “Basic” morphology parameters including : area, perimeter, height, width, major and minor axis lengths and angles, bounding box, centroid, solidity, circularity, aspect ratio, roundness |
| 14 | Compute convex hull area and perimeter | Useful for computing convexity indices |
| 15 | Save hull image | For rapid post-processing visual checks if needed |
| 16 | Redo a more stringent threshold to remove mixed background/plant pixels | The 2-3 pixels in the perimeter of the shape are a mix of background and plant color, and therefore need to be removed before measuring plant color parameters |
| 17 | Erode the borders of the plant to eliminate the edge pixels | To further remove mixed color pixels |
| 18 | Create a reduced mask based on the stringent threshold | To be applied on the RGB and HSB separated channels |
| 19 | Measure Red, Green, and Blue, densities on the reduced mask | The reduced mask is applied on each of the previously splitted R, G, and B channels. Measurements include mean density, stdev, mode, min, and max values |
| 20 | Measure Hue, Saturation, and Brightness densities on the reduced mask | The reduced mask is applied on each of the hue, saturation, and brightness channels. Measurements include mean density, stdev, mode, min, max, skewness, and kurtosis values |
| 21 | Export data to text file | All morphology and color measurements are saved in a csv file for further statistical analysis |

| Steps in Rstudio |  |  |
| --- | --- | --- |
| 22 | Extract metadata from the filenames | Get Plant id, camera id, date:time in separate fields |
| 23 | Merge image and plant data | Get Species, Room, and Location of each plant from a separate plant file |
| 24 | Compute days after sowing (DAS) and days under gradient conditions for each imaging time point |  |
| 25 | Perform visual quality check by plotting dimensions and color indices | For each species and time point, plotting Height vs Width indicates if there are clear abnormal measurements due to e.g. objects in the background. |
| 26 | Flag clear outliers | Outliers are flagged based on step 25 and on color measurements of the background reference card |
| 27 | Aggregate image data per plant | The measurements from the 6 side-view images are aggregated into one value per plant. For ex. Side area is the average of 6 images, side height and width are the maximum values.<br>Top- and side-view measurements are aggregated per plant and timepoint |
| 28 | Compute additional derived measurements | Voxel, Verticality, Green Leaf Index, Triangular Greenness Index, Chlorophyll content prediction are calculated |
| 29 | Merge imaging data and light mapping data | The local light data (intensity, spectra, computed Red:Blue ratio and phytochrome photostationary state (PSS)) is merged with plant imaging data |
| 30 | Merge imaging data and manual measurement data | E.g. leaf chlorophyll content recorded with Apogee probe |

**Table S2.** Summary list of the plant dimension, shape, and color parameters measured by imaging, including definition, calculation, and units.

| Label | Definition | Formula | Unit or scale |
| --- | --- | --- | --- |
| **Dimensions**<br>(note: For side-view parameters, the mean of the 6 images was used except for Height and Width where the max was considered more relevant) |  |  |  |
| HeightMax | Maximum height of the plant out of 6 side-view images during 180° rotation |  | mm |
| WidthMax | Maximum width of the plant out of 6 side-view images during 180° rotation |  | mm |
| AreaMean | Mean Projected Area out of 6 side-view images during 180° rotation |  | mm<sup>2</sup> |
| Area | Projected Area out of 1 top view image |  | mm<sup>2</sup> |
| MeanFeret | Average of maximum and minimum distances between 2 points along the selection boundary. |  | mm |
| Voxel | Plant volume estimate combining side- and top-view area of the plant | $\text{sqroot}(\max(\text{side-view area}) * \min(\text{side-view area}) * \text{top-view area})$ | mm<sup>3</sup> |
| **Shape factors**<br>(note: For side-view parameters, the mean of the 6 images was used) |  |  |  |
| Roundness | Degree of similarity to a circle derived from the fitted ellipse axes | minor axis / major axis (of the fitted ellipse) | Scale 0 to 1 |
| Solidity | Overall concavity derived from area and convex-hull measurements | area / convex-hull area | Scale 0 to 1 |
| Convexity | Edge "roughness" derived from convex hull and perimeter measurements | convex-hull perimeter / perimeter | Scale 0 to 1 |
| Circularity | Ratio of the area of the shape to the area of a circle having the same perimeter (a.k.a "isoperimetric quotient") | $4\pi * \text{area} / \text{perimeter}^2$ | Scale 0 to 1 |
| Compactness | Degree of compactness derived from the ratio of the diameter a circle with the same area to the major axis of the fitted ellipse | $\text{sqroot}((4/\pi) * \text{area}) / \text{major ellipse axis}$ | Scale 0 to 1 |
| **Color indices**<br>(note: For side-view parameters, the mean of the 6 images was used) |  |  |  |
| HueMean | Average hue component of the plant's color after transformation of the RGB image into HSB model (Hue Saturation Brightness) |  | Scale 0 to 255 |
| HueCv | Coefficient of variation of the plant's pixels hue | $\text{stdev}(\text{hue}) / \text{avg}(\text{hue})$ | % |
| SaturationMean | Average saturation component of the plant's color after transformation of the RGB image into HSB model (Hue Saturation Brightness) |  | Scale 0 to 255 |
| BrightnessMean | Average brightness component of the plant's color after transformation of the RGB image into HSB model (Hue Saturation Brightness) |  | Scale 0 to 255 |
| RedMean | Average red component of the plant's color in the RGB model |  | Scale 0 to 255 |
| GreenMean | Average green component of the plant's color in the RGB model |  | Scale 0 to 255 |
| BlueMean | Average blue component of the plant's color in the RGB model |  | Scale 0 to 255 |
| Density | Integrated density: The sum of the grey values of the pixels in the image or selection | $\text{area} * \text{mean grey value}$ | |
| GLI | Green Leaf Index : vegetation index for use with a digital RGB camera | $(2 * \text{Green} - \text{Red} - \text{Blue}) / (2 * \text{Green} + \text{Red} + \text{Blue})$ | |
| TGI | Triangular Greenness Index : approximate area of a triangle bounding a leaf reflectance spectrum, where the vertices are in the red, green, and blue wavelengths. | $((670 - 480) * (\text{tRed} - \text{tGreen}) - (670 - 550) * (\text{tRed} - \text{tBlue})) / -200$ | |
| Chl_predicted | Predicted leaf chlorophyll content derived from multiple linear regression using Red Green and Blue components of the plant color in the RGB model | $440 + \text{blue} * 7.266 + \text{red} * 10.873 + \text{green} * -15.545$ | $\mu\text{moles/m}^2$ |

**Figure S1.**
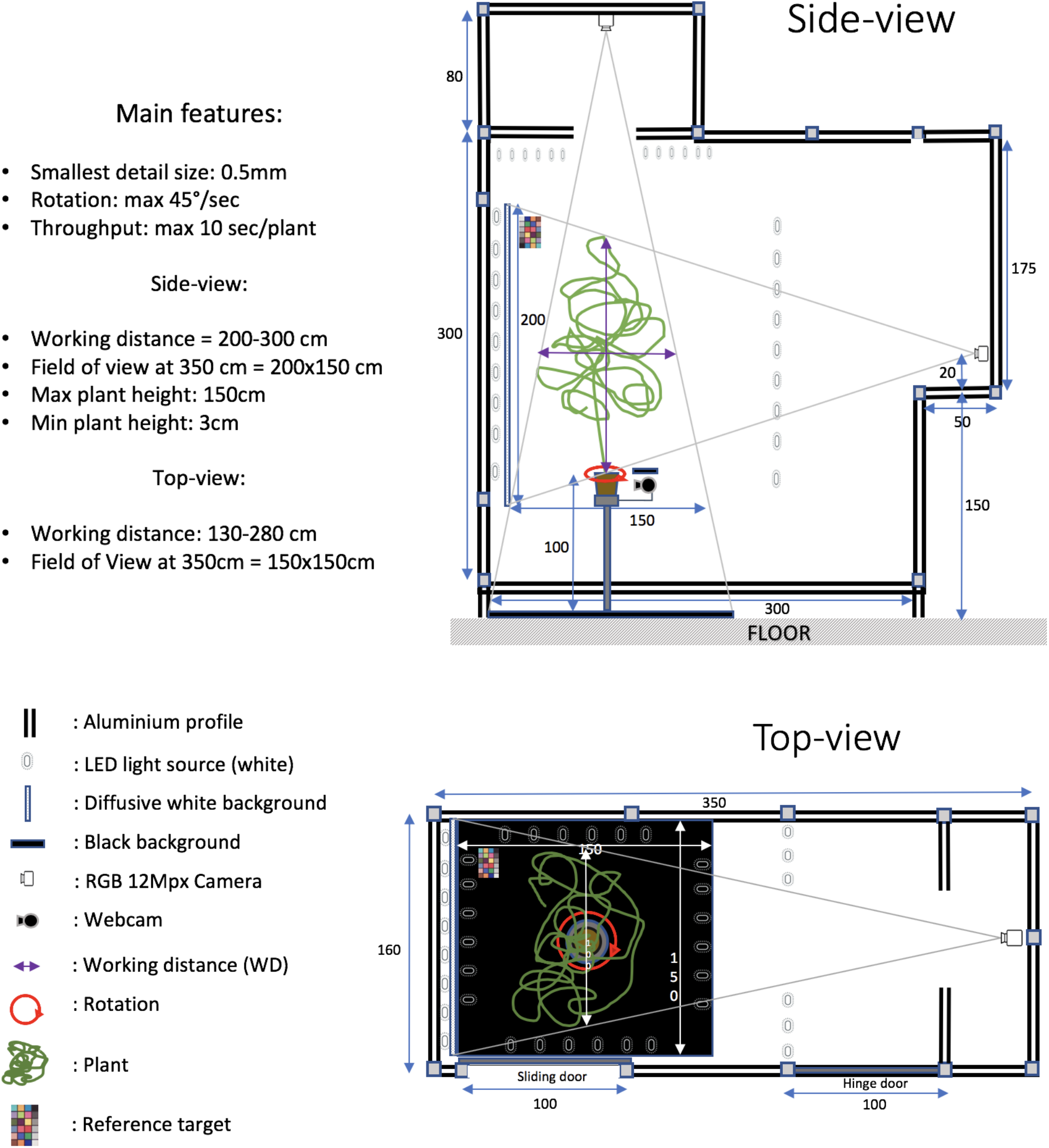
Schematic representation of the imaging setup.

**Figure S2.**
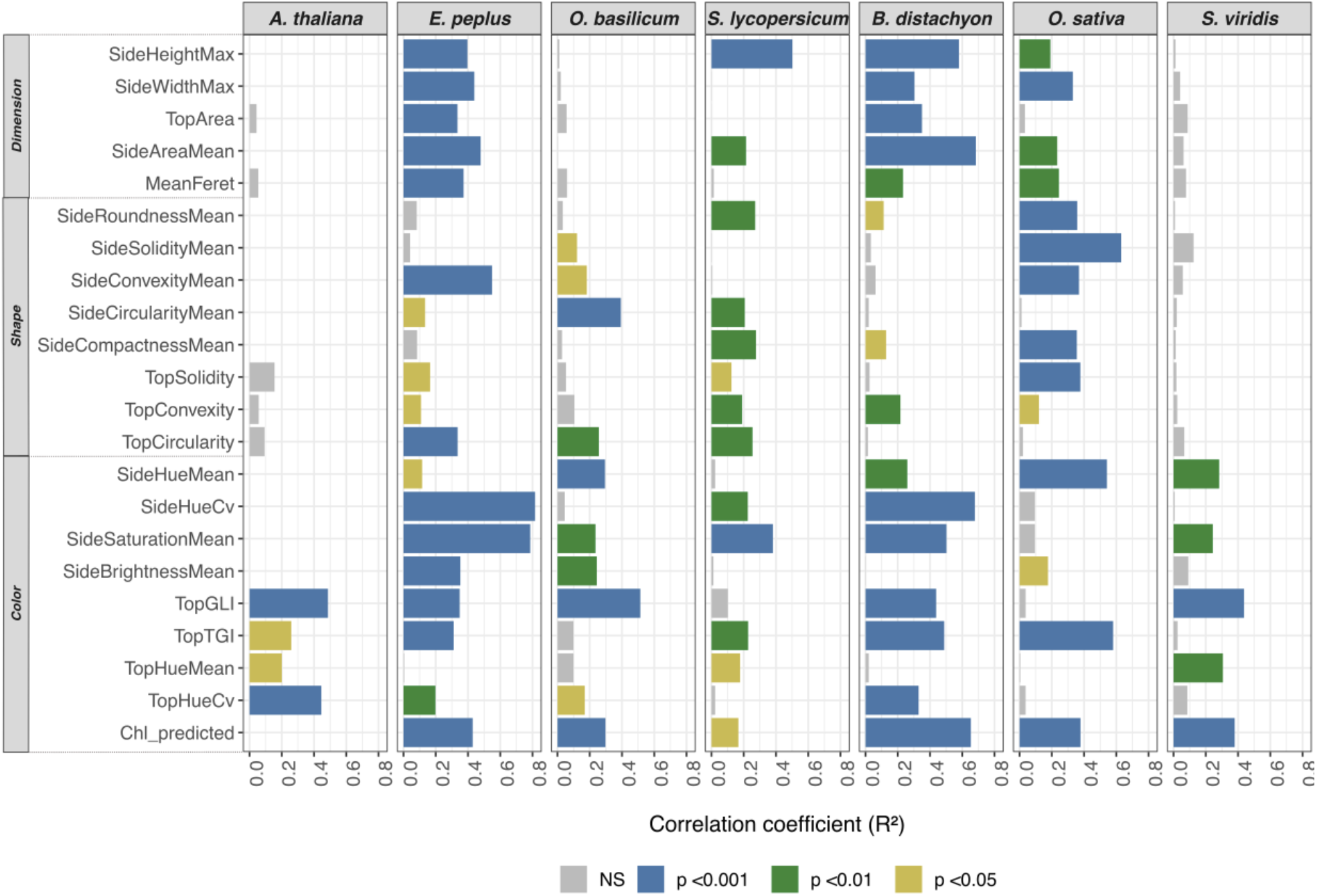
Correlation coefficients (R^2^) of the linear regressions between various phenotypic traits measured at day 29 after the start of the gradient treatment and the Red:Blue ratio, as shown in Figure 6.

**Figure S3.**
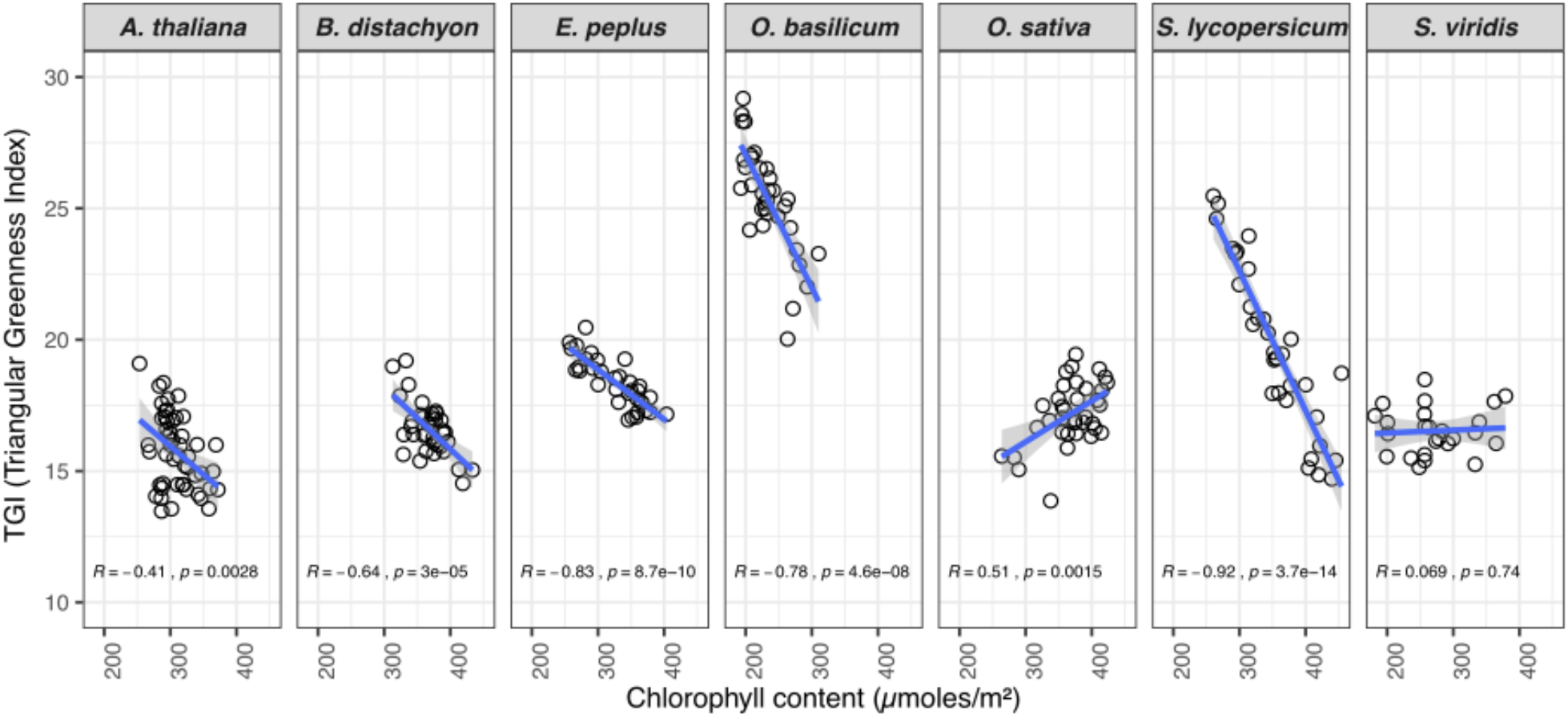
Correlation between leaf chlorophyll content, as measured manually with an Apogee MC-100 chlorophyll meter, and the Triangular Greenness Index (TGI) computed from RGB images.

